## Supplemental figures for "Increased inflammatory signature in myeloid cells of non-small cell lung cancer patients with high clonal hematopoiesis burden"

**Running title:** Increased inflammatory signature in CHIP-positive NSCLC patient blood

**Corresponding author:**

Murim Choi, Department of Biomedical Sciences, Seoul National University College of Medicine,103 Daehak-ro, Jongno-gu, Seoul 03080, Republic of Korea

Se-Hoon Lee, Division of Hematology-Oncology, Department of Medicine, Samsung Medical Center, Sungkyunkwan University School of Medicine, 81 Irwon-ro, Gangnam-gu, Seoul 06351, Republic of Korea

**Supplementary Tables supplied in a separate Excel file**

**Supplementary Table S1**. Clinical information of NSCLC cohort with ICI treatment.

**Supplementary Table S2**. Information of control samples.

**Supplementary Table S3**. List of CHIP variants detected from targeted sequencing.

**Supplementary Table S4**. List of CHIP variants detected from WES.

**Supplementary Table S5**. Identified DEGs from annotated clusters of scRNA-seq.

**Supplementary Table S6**. Identified DEGs from each cluster by VAF bin.

**Supplementary Table S7.** Gene-Module annotation from WGCNA analysis

**Supplementary Figures**

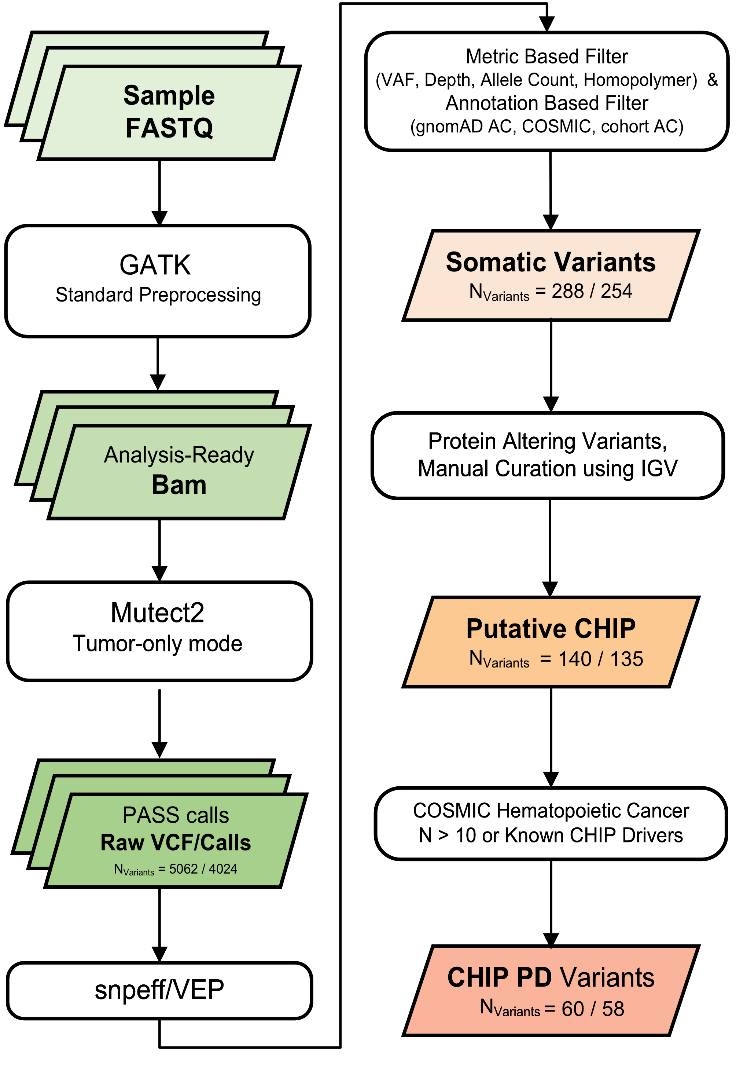

**Supplementary Fig. S1** Variant calling and filtering scheme. We collected Numbers indicate count of variants after each variant filtering scheme **(all samples/excluding controls).**

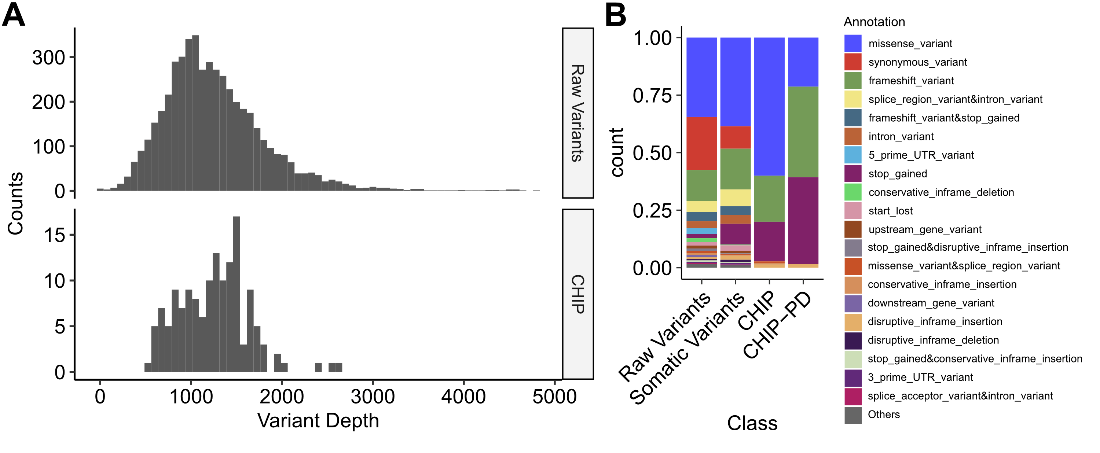

**Supplementary Fig. S2** Depth and annotation distributions from variants filtering scheme.

**A** Variant depth distributions from raw calls and putative CHIP variants. **B** Distribution of variant annotation from each variant filtering step.

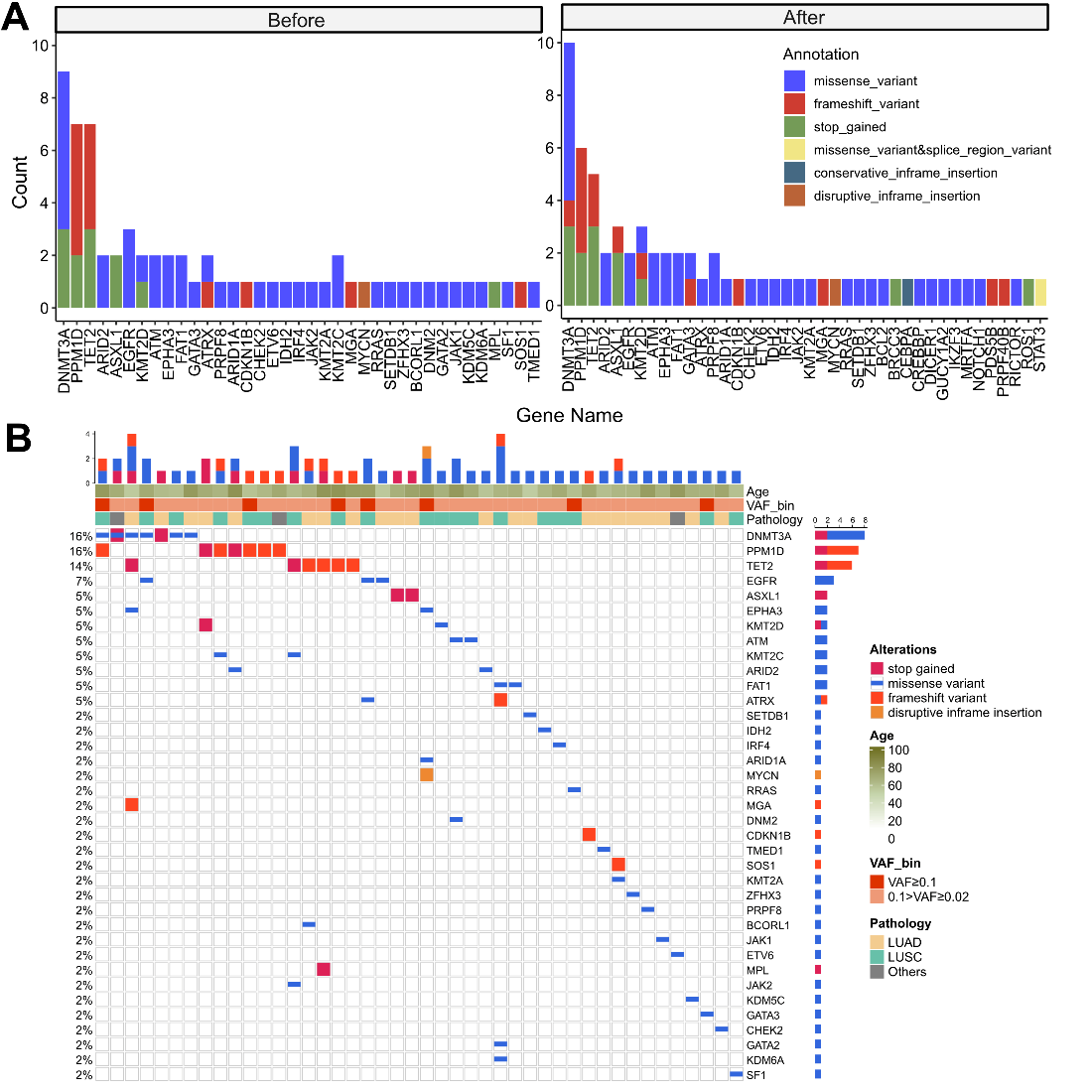

**Supplementary Fig. S3** CHIP profiles from discovery cohort with panel sequencing (sample *n* = 100/91 for before treatment and after treatment). **A** Gene distribution in before- and after- ICI treatment. Color indicates variant annotation. **B** Waterfall plot showing CHIP mutations from baseline, cohort samples.

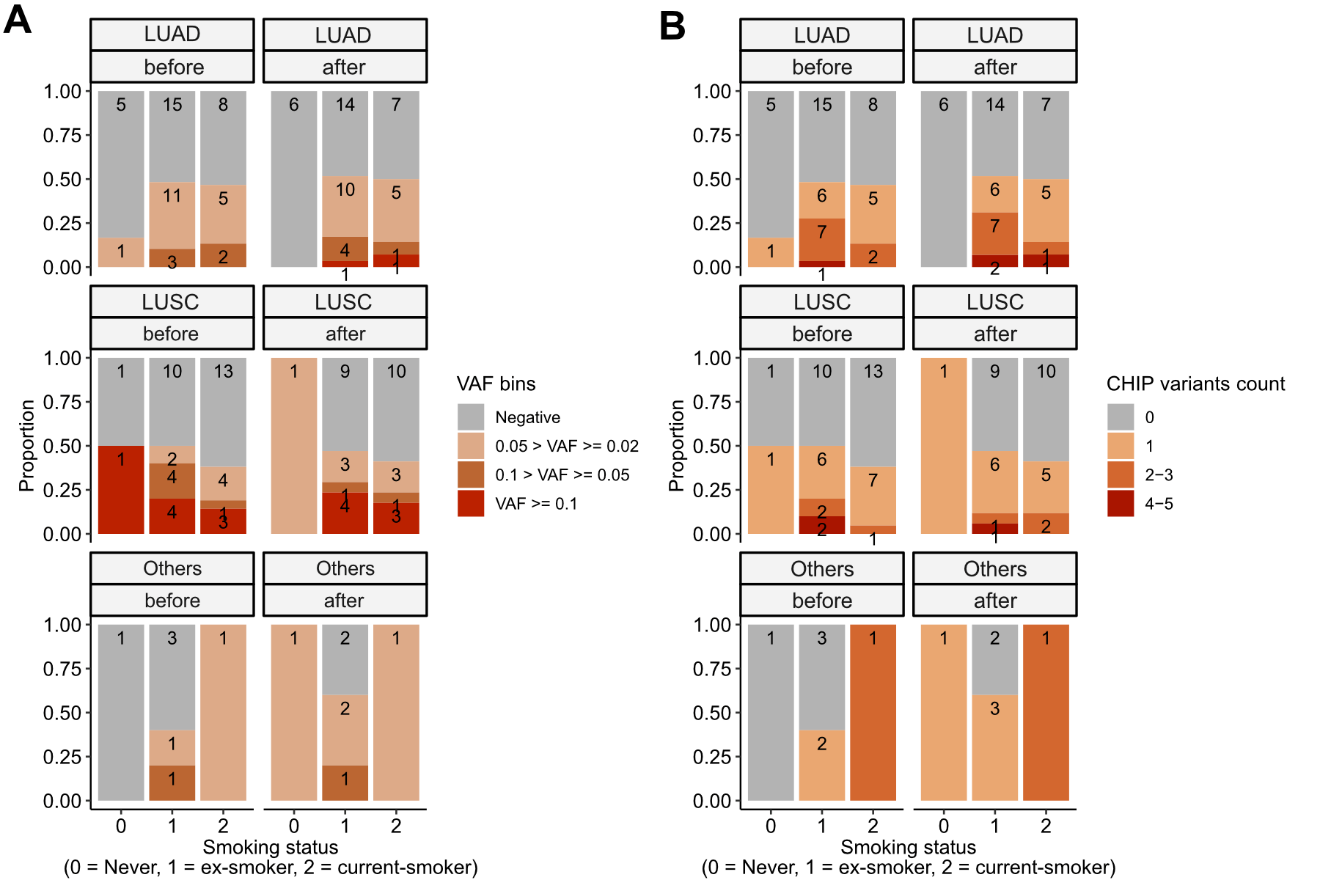

**Supplementary Fig. S4** CHIP profiles, by stratified smoking status. CHIP status was displayed using **A** binned VAF and **B** variant count per patient. Colors indicate binned VAF and variants count.

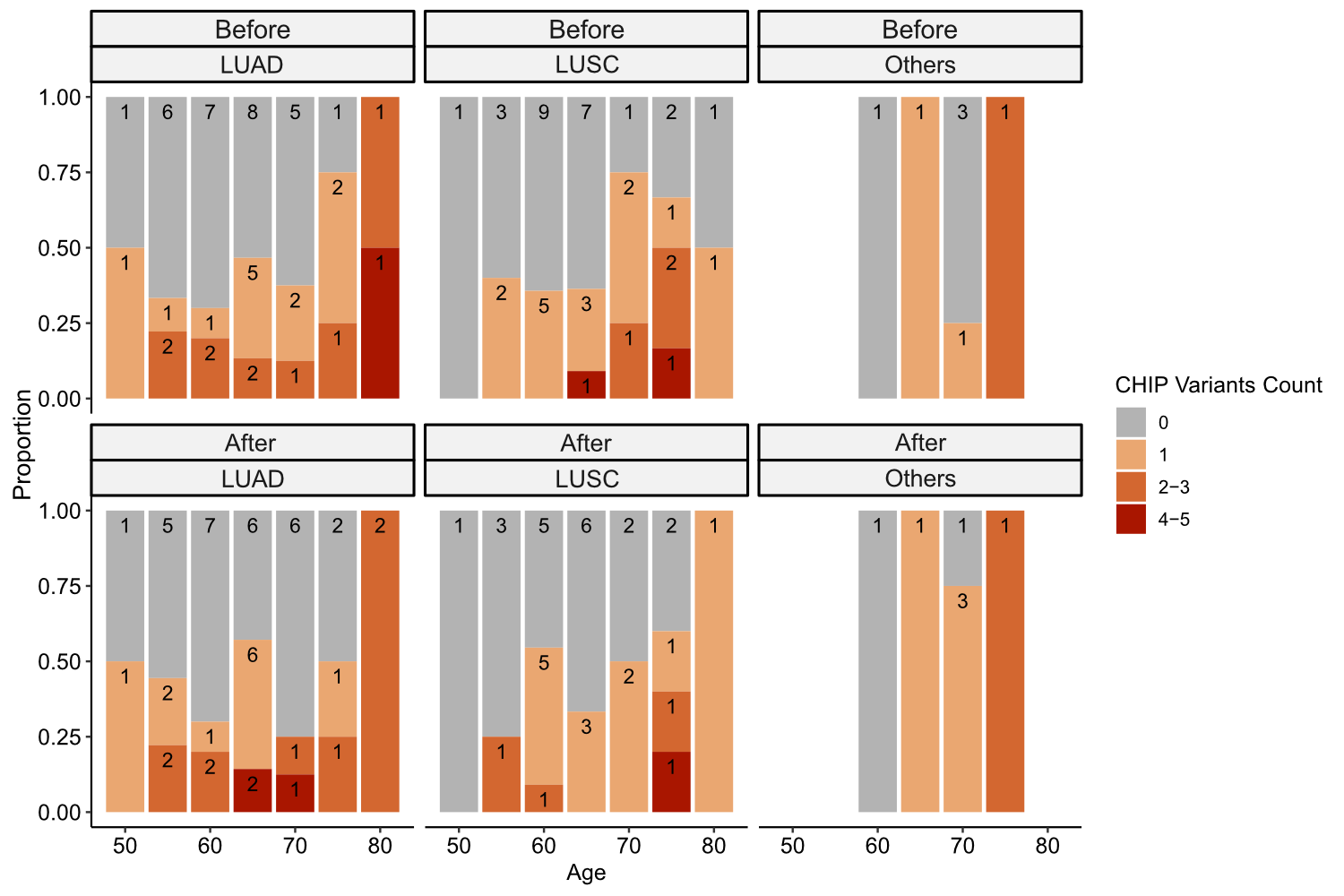

**Supplementary Fig. S5** Relationship between CHIP variants counts and clinical parameters. CHIP variants counts and major clinical parameters were presented. Color indicate the binned number of CHIP variants.

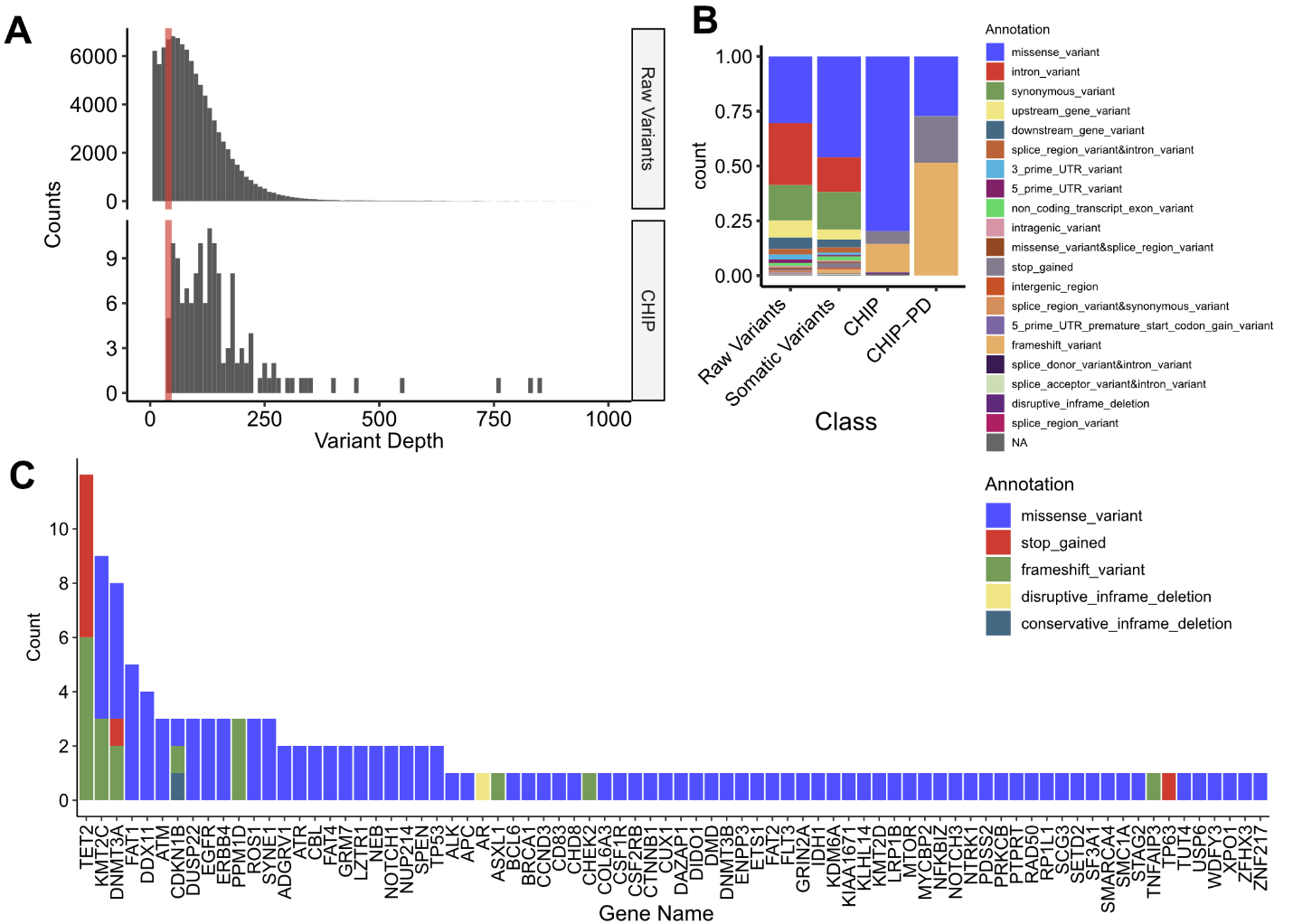

**Supplementary Fig. S6** CHIP profiles from replicative cohort with PBMC WES (*n* = 180).

**A** Variant depth distribution from raw and CHIP variant. Colored Bar indicates variants cutoff, with depth = 40. **B** Distribution of variant annotation from each variant filtering step. **C** All genes from CHIP variants. Colors denote variant annotation.
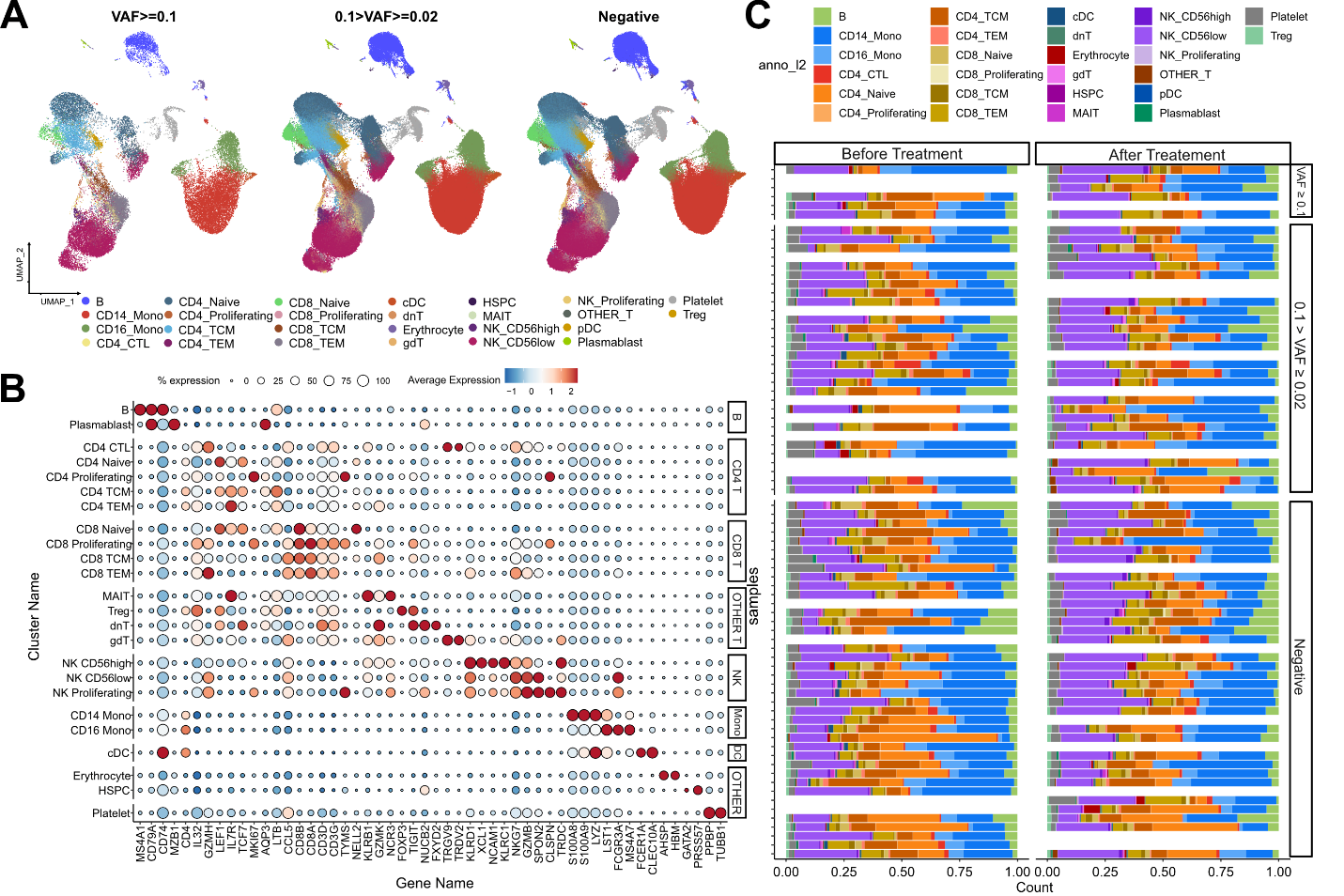
**Supplementary Fig. S7** scRNA-seq profile of ICI-treated, NSCLC patients. **A** UMAP plot of scRNA-seq data, with CH VAF bins. **B** Cell-type specific marker gene expression from each clusters. **C** Cell composition plot from each sample.

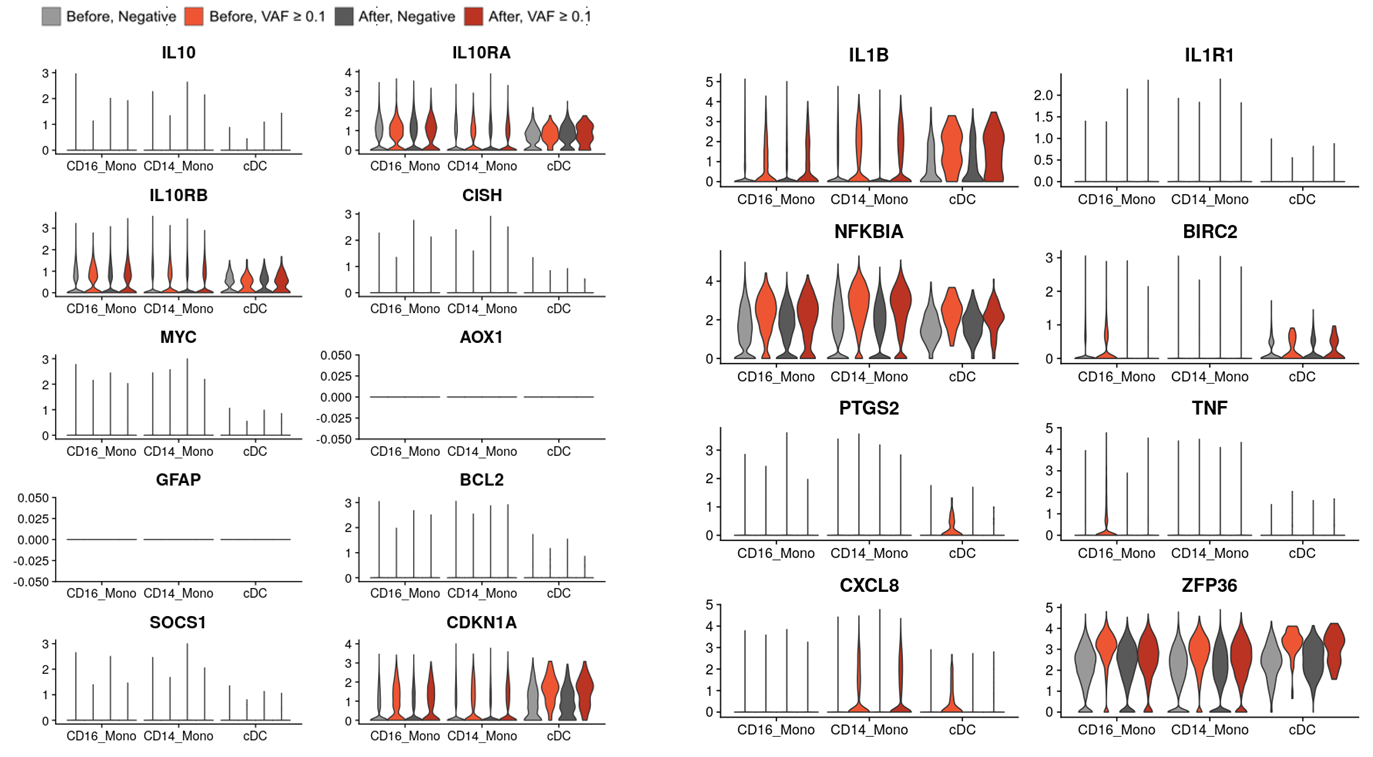

**Supplementary Fig. S8** Gene expression of selected pathways for **A** IL-10 and **B** IL-1B.

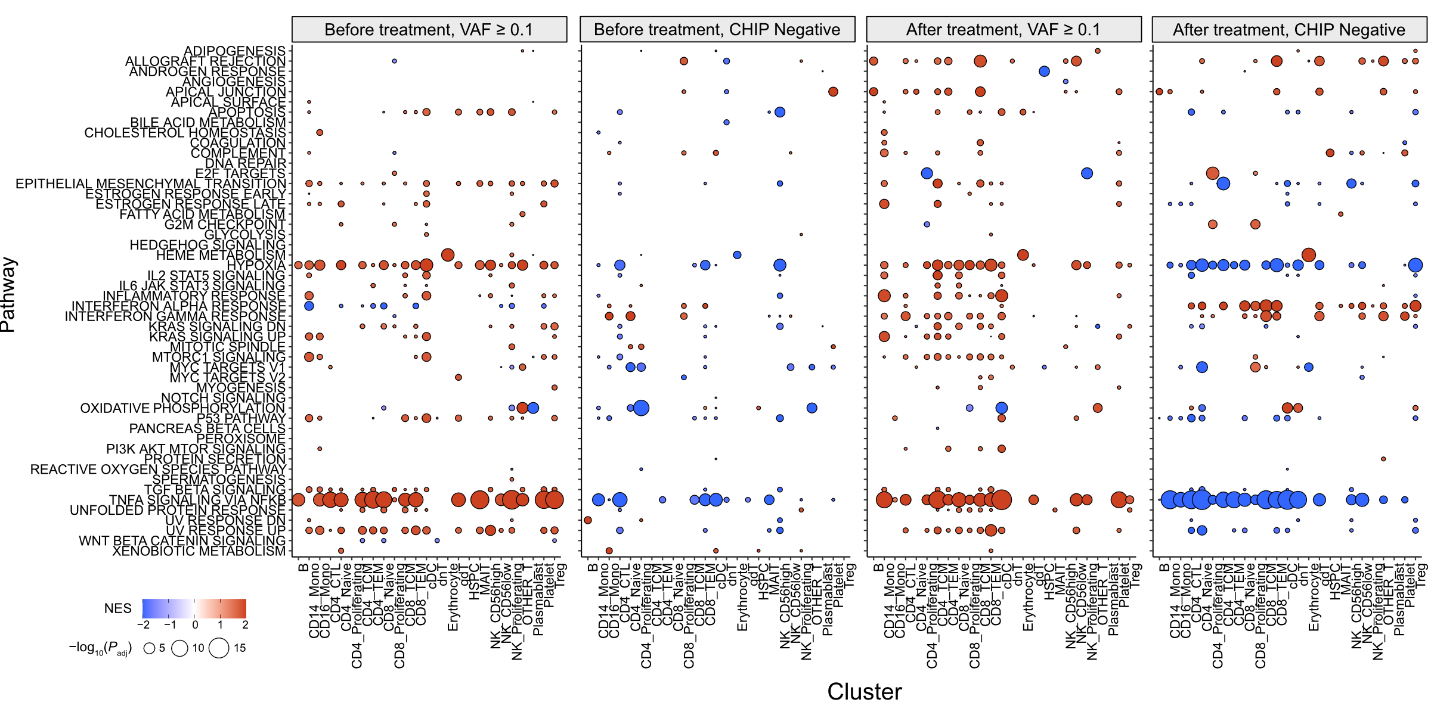

**Supplementary Fig. S9** GSEA analysis results from all annotated clusters.

Color represents normalized effect score (NES), and dot size represents adjusted *P*-values.

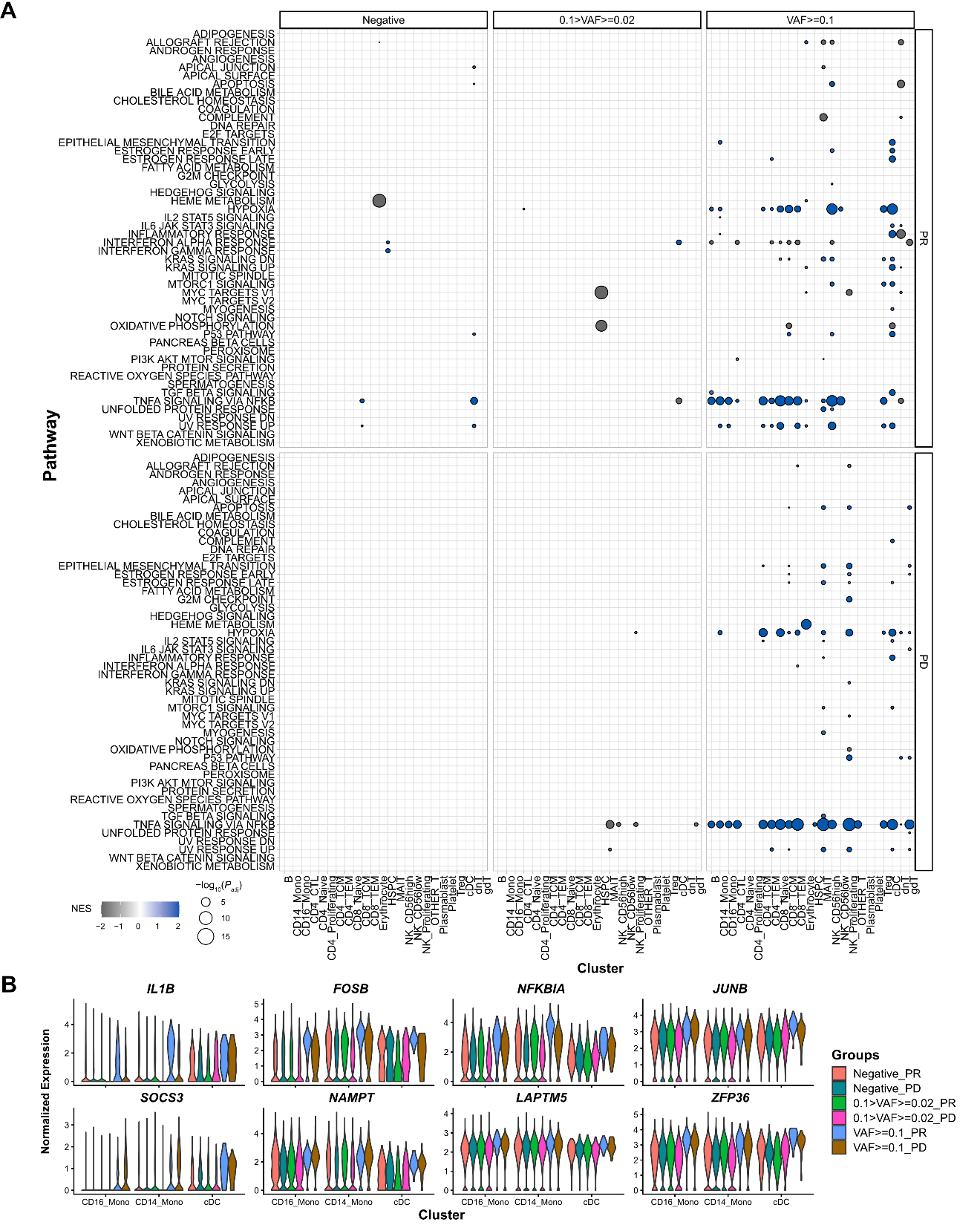

**Supplementary Fig. S10** GSEA results and representative gene expression patterns, associated with ICI response and CHIP VAF bin. **A** Results of GSEA analysis in response to ICI and CHIP VAF bins, from before treatment samples. Color represents normalized effect score (NES), and dot size represents adjusted *P*-values. **B** Representative gene expression is displayed using same panels as **Fig. 3B**. Colors indicate groups based on CHIP VAF bin and ICI response status.

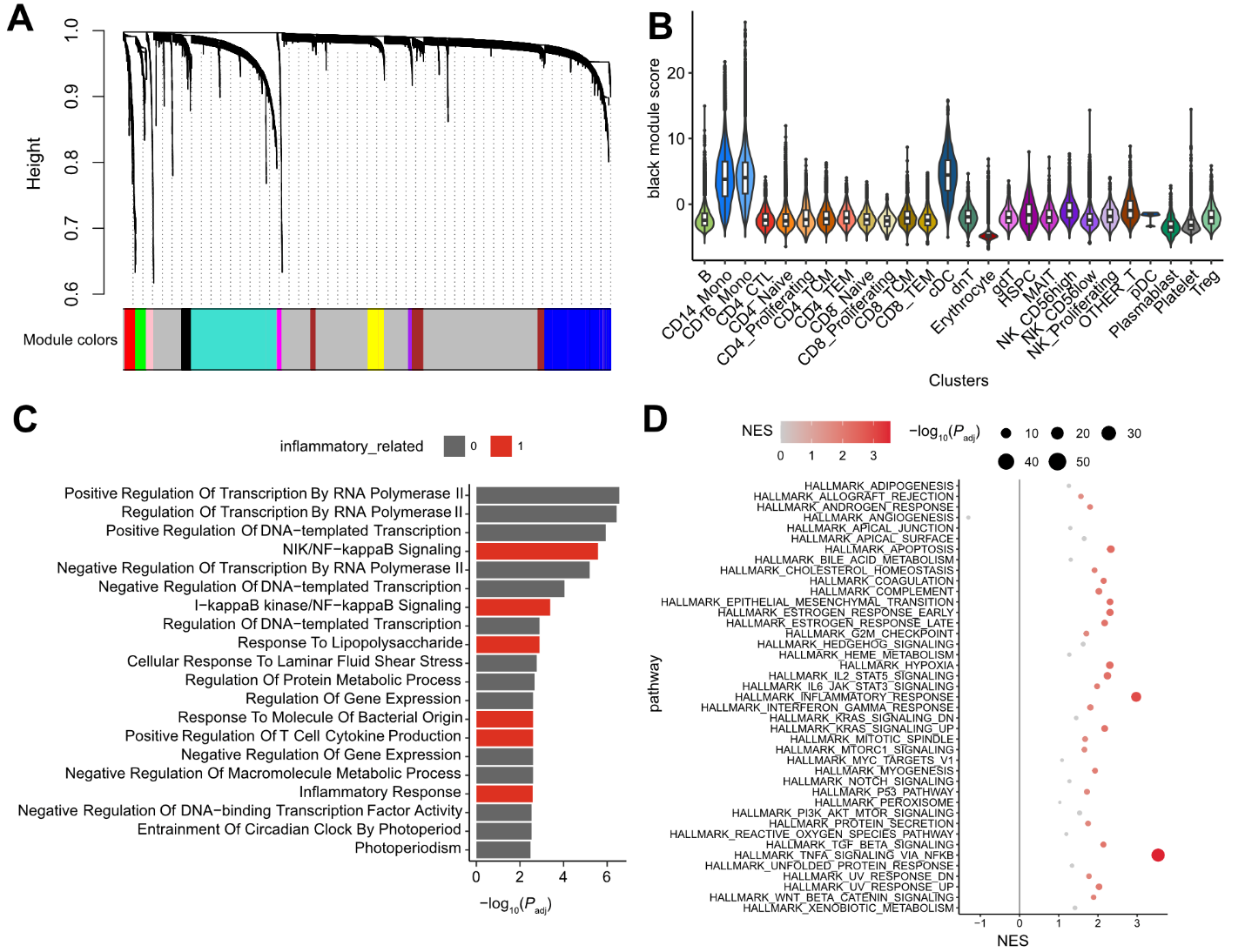

**Supplementary Fig. S11** scRNA-seq WGCNA analysis using hdWGCNA. **A** WGCNA dendrogram, derived from myeloid gene expressions. **B** cluster-wise ‘black’ module score distribution. **C** GO enrichment analysis of the Black module genes. The colored bar indicates inflammatory pathway-related GO terms with significant enrichment. **D** GSEA analysis of Black Module Genes. Colored dots indicate significant gene set enrichment.

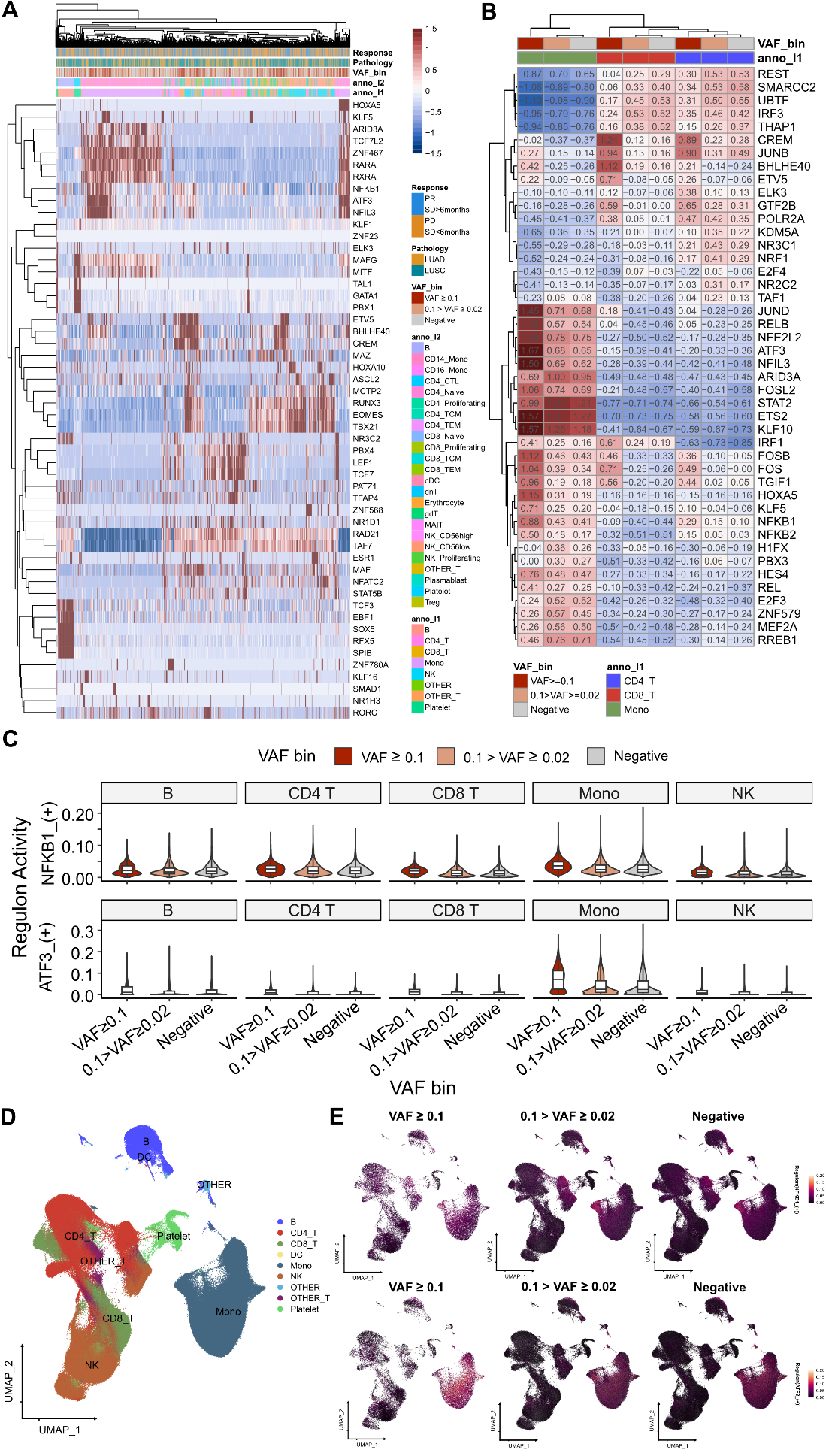

**Supplementary Fig. S12** SCENIC analysis for gene regulatory networks (GRN) on scRNA-seq. **A** Clustered, representative GRNs from scRNA-seq dataset. 1,500 cells from each VAF bin were presented for this visualization. Color represents GRN AUC scores. **B** Aggregated GRN scores from cluster and VAF bins. **C** NFKB1 and ATF3 regulon scores in major cell clusters, stratified by VAF bins. **D** UMAP plot for clusters, used in SCENIC analysis. **E** Regulon expression distribution, using Seurat’s *featureplot.*

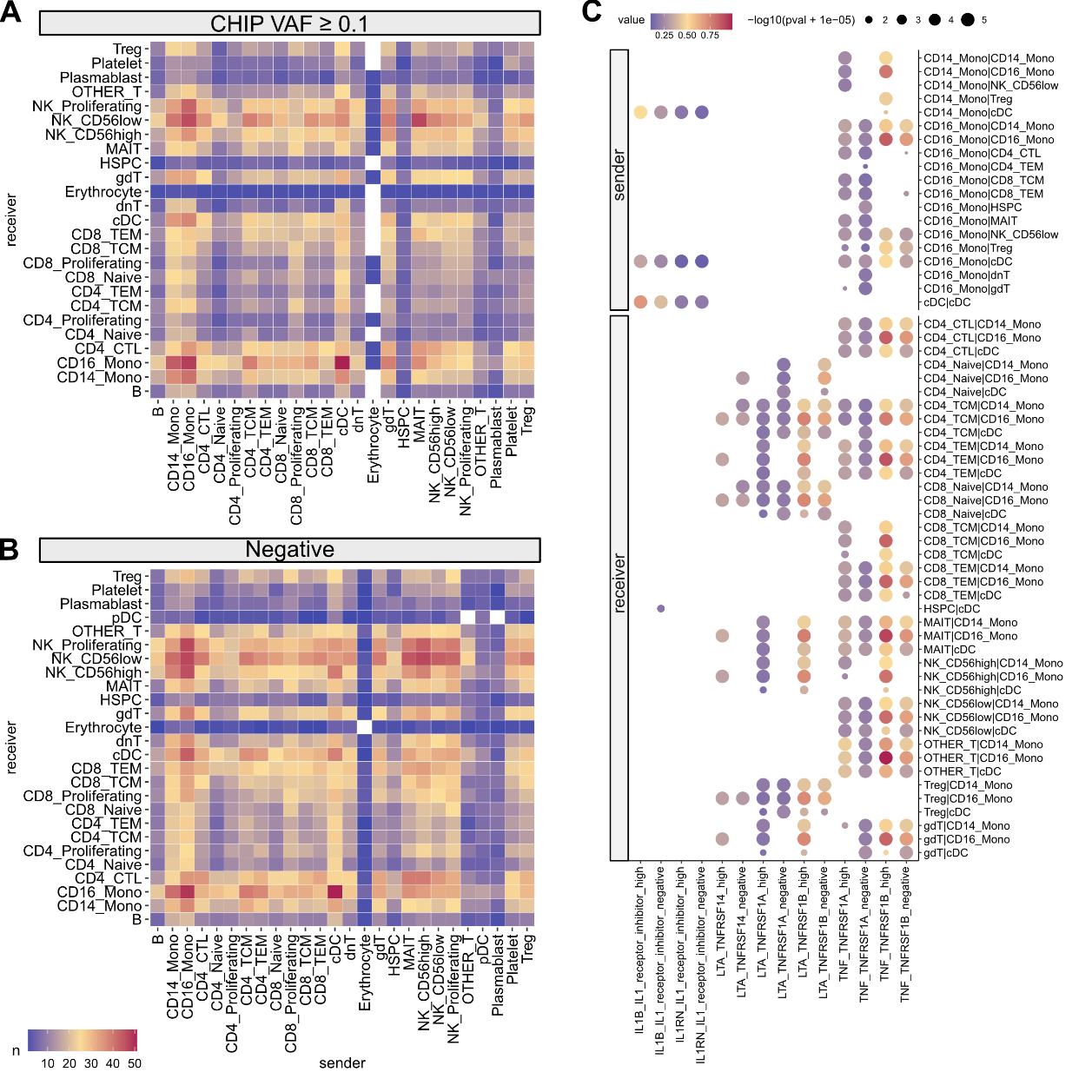
**Supplementary Fig. S13** Cell-cell interaction analysis using CellphoneDB. Heatmap illustrating the numbers of cell-cell interactions using **A** high VAF CHIP and **B** CHIP negative samples. **C** Selected interaction pathways related to myeloid cells are displayed. Dot sizes represent interaction p-values, and colors indicate the magnitude of interactions. In row names, the order of names is presented as sender | receiver.

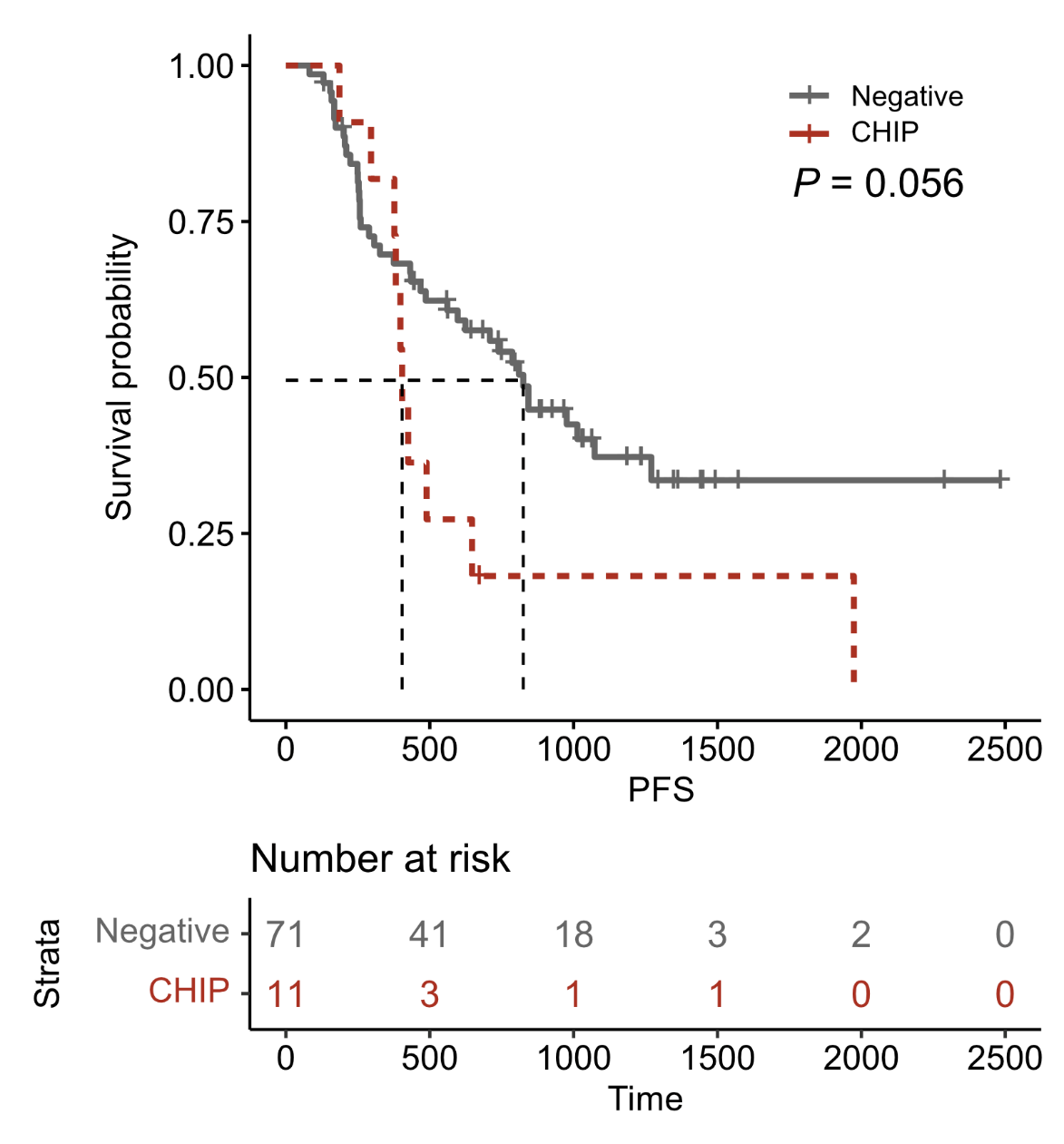

**Supplementary Fig. S14** Survival plot from this cohort. Survival curve using progression-free survival in CHIP Negative (N = 71) and high-burden CHIP (N = 11), using *survplot* in R.

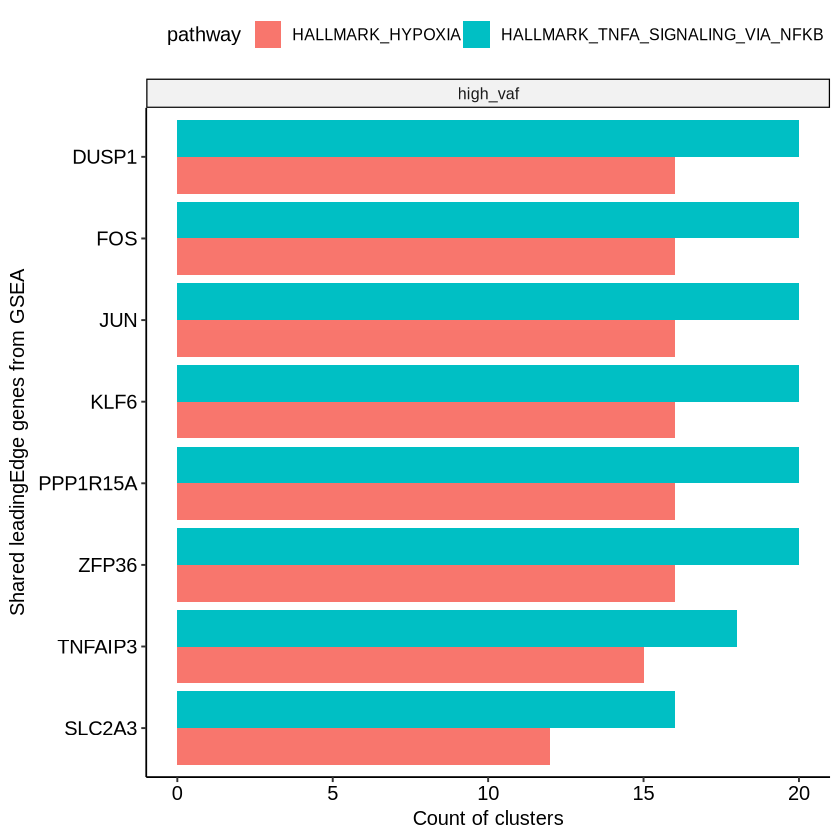

**Supplementary Fig. S15**: Bar plot of the top shared GSEA leading edge genes, using significantly enriched (adjusted P < 0.05 for each cluster) Hypoxia and TNFɑ signaling pathways across clusters. The bars represent the count of clusters in which each gene is included in the leading edge.
